# Optimizing storage, high-molecular weight DNA extraction and genome reconstructions from human faecal samples

**DOI:** 10.1101/2025.08.27.672560

**Authors:** Klara Cerk, Anthony Duncan, Nabina Shrestha, David Baker, Muktha Kukreja, Jennifer Ahn-Jarvis, Rob James, Christopher Quince, Falk Hildebrand

## Abstract

Metagenomic sequencing has become an essential tool for assessing the composition and functional potential of microbial communities. However, established metagenomic DNA extraction protocols are limited by trade-offs between universal taxonomic representation, DNA quality, quantity and molecule length, making them mostly unsuitable for 3^rd^ generation sequencing.

**Methods:** We assessed the effectiveness of high molecular weight (HMW) DNA workflows from faecal samples, using different combinations of collection, storage and extraction kits. Faecal samples were either immediately processed, frozen at -80°C or stored in DNA/RNA Shield, GutAlive or OMNIgeneGUT and then subjected to three different DNA isolation kits, each modified for mechanical, chemical and enzymatic cell lysis steps, totalling 32 evaluated combinations. Isolated DNA was assessed for its quality and quantity, while taxonomic consistency was evaluated using metataxonomics and validated with metagenomic sequencing.

**Results:** The yield of HMW DNA differed substantially between storage and DNA extraction protocols. Specifically, storage protocol had the highest impact on HMW DNA recovery, with DNA/RNA Shield storage yielding on average 51% more HMW DNA >30kb. Similarly, the sample storage had independently the effect size of *R*^2^ = 26.9% on the microbiome composition, and since the DNA extraction protocol in our study inherently combines both DNA extraction kit and lysis type, the latter had the largest effect size (*R*^2^ = 12.8%).

While all kits had specific dis/advantages and biases, DNA/RNA Shield and Maxwell RSC PureFood GMO and Authentication kit performed best overall for long-read metagenomics. The protocol was validated on a) isolate genomes and b) 4 faecal samples, demonstrating both increased DNA yield at >30kb molecule length. Using the MG-TK pipeline these metagenomes yielded genomes on average, per sample, 48 Pacific Biosciences (PacBio) and 55 Oxford Nanopore Technologies (ONT) circular bacterial metagenomic assembled genomes.

**Conclusion:** The current gold-standard of freezing faecal samples was surprisingly inconsistent and inefficient for long-read metagenomics, necessitating rethinking gut microbial study designs. We provide a simple and cheap protocol that preserves both taxonomic composition and HMW DNA quality, while our bioinformatics workflow allows for complete bacterial genome reconstruction.

## Background

Metagenomic sequencing has revolutionized our understanding of microbial communities, enabled through direct sequencing of DNA from a sample. This has led to important breakthroughs, such as the creation of microbial gene catalogues (*1*, *2*), recovery of microbial genomes directly from metagenomes [3, 4] and strain-resolved comparisons of different microbial strains found between samples [5]. Up to today, most metagenomes are sequenced using short-read sequencing (typically 150-300 bp in length). However, rapid advances in long-read sequencing technologies now offer novel opportunities for high-precision metagenomics, such as a characterisation of long continuous DNA molecules that span genomic repeats and structural variants otherwise difficult to resolve with short-read sequencing [6].

These advances are counterbalanced by long-read sequencing being challenging to use in a metagenomic context. This is because the most common metagenomic DNA extraction kits are designed for short-read sequencing [7], relying on bead-beating for cell lysis, and spin-columns for nucleic acids purification (Suppl. Table 1). Long-read sequencing generally requires high concentrations of highly purified DNA of high molecular weight [8], which limits the utility of established DNA extraction kits designed for short-read sequencing. Specifically bead-beating, a physical disruption of biological cell membranes, has proven essential to obtain taxonomically unbiased representations of DNA from samples of different environments [9], but can be detrimental to the recovery of long DNA molecules particularly in combination with spin columns [10, 11], as the mechanical forces involved result in DNA shearing which can lead to lower yields of HMW DNA.

For third generation sequencing, it is imperative to obtain HMW DNA, ideally in the range of 7-15 kb for PacBio and >40kb for ONT sequencing. Although several specialized kits are now available for extracting HMW DNA (Suppl. Table 1), these rely exclusively on gentler enzymatic or chemical lysis steps [7, 11, 12]. This is problematic for metagenomic sequencing, as this likely introduces taxonomic biases, e.g. the ratio of easy (Gram-negative) to hard (Gram-positive) to lyse bacterial cells are of concern when using chemical and enzymatic lysis [8, 13].

Accordingly, sample storage protocols that ensures the short- and long-term preservation of microbial diversity is crucial to conduct cost-efficient clinical trials [14]. The immediate freezing and storage of samples at the point of collection (−20 to -80 °C) has been widely accepted as best practice for sequence-based analyses [15]. However, recent advancements involve storing samples with chemical stabilizers, such as DNA/RNA Shield or OMNIgeneGUT, which inhibit bacterial growth and may be more practical for both experimental and the clinical setting then freezing samples [16]. These preservatives which contain chelating agents, can stabilize the nucleic acids contained in a sample at ambient temperatures, allowing storage for more than a year, while preserving the taxonomic and (gene) functional composition. New protocols are required to take advantage of these technological advances, spanning from sample preservation and DNA extraction to bioinformatic processing.

Here we tested the combined effect of different faecal sample storage and DNA extraction methods in their effectiveness at obtaining HMW DNA, quantifying total yield, concentration, purity, integrity and suitability for long-read sequencing. We found that sample storage had greater impact on DNA length and yield recovery than different extraction protocols or lysis steps. The recommended combination of methods was further validated using mock communities and single isolates, as well as on stool samples using Illumina, ONT and PacBio sequencing. Using our recommended bioinformatic workflow, this enabled taxonomic robust community profiling and the recovery of completely circular bacterial genomes.

## Materials and methods

### Storage and sample collection methods

The Quadram Institute of Bioscience (QIB) HNU**/**CRF facility collected stool samples from two QIB Participant / Donor NRP biorepository participant’s database (“V1”, “V2”) and handled under ethics approval UEA FMH ref: 201414-06HT and NNUH R&D ref: 70-05-19. Each participant received a collection package containing a biodegradable “Faeces Catcher”, disposable gloves, an airtight pouch, an anaerobic pouch AnaeroGenTM (ThermoFisher Scientific), absorbent sheets, and a blue cooling box, along with instructions for the collection procedure. After participants collected their samples, these were packaged with anaerobic sachets and kept in cooled conditions at approximately 4°C to prevent degradation or contamination during delivery to laboratory, usually within 30 minutes of defecation.

The consistency of faecal samples was assessed on Bristol Stool chart before being homogenized and subsequently divided into four parts to test different storage conditions: Immediate processing (“I”): Approximately 200mg of stool sample was aliquoted into replicates from which DNA was immediately extracted using different extraction protocols, to establish baseline controls.

Frozen stool samples (“F”): Stool was aliquoted into 500mg portions that were immediately frozen at -80°C and stored for 4 days prior to processing and extraction.

The DNA/RNA Shield Stabilization Solution (Zymo Research International, R1100) (“S”): A spoonful of stool sample (1g) was placed in the collection tube containing 4.5 mL of preservation liquid and 5 glass beads (Zymo Research International, R1101). For stool sample V2 we also tested different amount of a faeces sample, summarized in Suppl. Table 2. The sample was then homogenised by manual shaking to create a suspension as per the manufacturer’s instruction. Aliquots were kept at room temperature (RT, 21°C) or at 30°C for 4 days, before being processed.

The GutAlive Anaerobic Microbiome Collection Kit (MicroViable Therapeutics SL, 7245-PS) (“G”): the faecal sample was split into two separate collection devices, the collection tube prepared as per manufacturer’s instruction and kept at either RT or 30°C.

In addition, for stool sample V2, we also tested:

The OMNIgeneGUT Stabilization Solution (DNA Genotek, OMR-200) (“O”): as per manufacturer’s instruction, 500mg of stool sample was scooped into collection tube containing 2 mL of preservation liquid and one glass bead and thoroughly shaken to create a suspension. Aliquots were kept at RT and at 30°C. Detailed storage conditions can be found in Suppl. Table 2.

### DNA Extraction kits and cell lysis type

Three separate extraction kits were combined with up to four alterations in protocol use, namely cell lysis type. The combination of DNA extraction kit and lysis type was combined in variable “Protocol”. The commercial kits used were Maxwell PureFood GMO and Authentication kit (Promega, AS1600) (“MPureFood”), Maxwell RSC Fecal Microbiome kit (Promega, AS1700) (“MFaecal”), and the QIAmp PowerFaceal pro kit (QIAGEN, 51840) (“QIAFeceal”).

Up to four variants of each kit were used a subset of kit-provided reagents to achieve microbial cell lysis via: i) chemical, ii) lytic enzymes (enzymatic digestion), iii) bead-beating via Lysis Matrix E (MP Biomedicals), or iv) bead-beating via PowerBead Tubes, Ceramic 1.4 mm (QIAGEN). Bead beating was conducted using Vortex adapter for 24x2ml tubes for Vortex-Genie™2 (10 min, highest speed) (Scientific Industries SI™), as well as FastPrep-24™ (45s at 6.0 m/s) (MP Biomedicals). After cell lysis, subsequent purification was performed with the use of DNA binding columns (with QIAFeceal kit) or use of DNA binding magnetic beads (with MPureFood and MFaecal), as provided in each kit. Protocols and alterations are summarized in Suppl. Table 2 and Suppl. Table 3.

Duplicates were made for each extraction technique to ensure reproducibility and aid in the validation of results.

### Assessment of selected protocol on single isolates and mock community

To validate the optimal protocol and storage method identified in stool (i.e., Protocol 3 with DNA/RNA Shield preservation), we tested its performance on pure bacterial cultures and a defined mock community. Six bacterial species were cultured in LB broth for 24 h at 37 °C: two Gram positive (*Streptococcus dysgalactiae*, *Staphylococcus epidermidis*) and four Gram negative (*Escherichia coli*, *Citrobacter freundii*, *Enterobacter aerogenes*, *Salmonella enterica*). *E. coli* and *S. dysgalactiae* were also processed as single isolates, while the mock community contained all six strains mixed in equal proportions. For each sample, 1 mL of culture was pelleted (9,000×g, 1 min), resuspended in 1 mL DNA/RNA Shield Stabilization Solution (Zymo Research, R1100), and stored for 24 h at -20°C prior to extraction. DNA was extracted using the Protocol 3 i.e., bead-beating via Lysis Matrix E (MP Biomedicals) and Vortex adapter for 24x2ml tubes for Vortex-Genie™2 (10 min, highest speed) (Scientific Industries SI™) and the Maxwell PureFood GMO and Authentication Kit (Promega, AS1600).

### Assessment of DNA quality and quantity

Extracted DNA quantity was measured using the Invitrogen Qubit 4 Fluorometer (Thermo Fisher Scientific, Waltham, USA) with Qubit dsDNA HS Assay Kit (Thermo Fisher Scientific, Waltham, USA) from 2 µl of DNA extract. The purity of DNA eluates were evaluated by measuring absorbance at 230 nm, 260 nm, and 280 nm with the NanoDrop 2000 Spectrophotometer (Thermo Fisher Scientific, Waltham, USA). Pure DNA has a 260/280 ratio around 1.8, and a 260/230 ratio that ranges between 2.0 and 2.2. The average fragment length of the extracted DNA was determined by using the 4200 TapeStation System (Agilent, Santa Clara, USA) with the Genomic DNA ScreenTape and reagents (Agilent, Santa Clara, USA).

### 16S rRNA gene amplification, library preparation and sequencing

Extracted genomic DNA was normalised to 5ng/µl with EB (10mM Tris-HCl). A PCR master mix was made up using 10 µl KAPA 2G Fast Hot Start Ready Mix (Merck), 0.1 µl 100 µM forward tailed specific primer, 0.1 µl 100 µM reverse tailed specific primer [17] and 7.8 µl PCR grade water per sample. 18 µl master mix were added to each well to be used in a 96-well plate followed by 2 µl of DNA and mixed. Specific PCR conditions were 95°C for 5 minutes, 30 cycles of 95°C for 30s, 55°C for 30s and 72°C for 30 seconds followed by a final 72°C for 5 minutes. For the second PCR 10 µl KAPA 2G Fast Hot Start Ready Mix (Merck) and 8µl PCR grade water were mixed per sample and added to a 96 well plate. 1µl of 10µM 8bp Unique Dual Indexes were added to each well. Finally, 1 µl of PCR 1 was transferred into the PCR 2 master mix plate. The second PCR was run using 95°C for 5 minutes, 10 cycles of 95°C for 30s, 55°C for 30s and 72°C for 30 seconds followed by a final 72°C for 5 minutes. Final libraries were quantified by Qubit and equimolar pooled together. A single 0.7X SPRI clean-up using sample purification beads (Illumina® DNA Prep, (M) Tagmentation (96 Samples, IPB)) was performed on the pool. After the final quality check using Qubit and Tapestation, the pool was run at a final concentration of 750pM on an Illumina Nextseq 2000 instrument using a NextSeq 2000 P1 XLEAP-SBS Reagent Kit (600 Cycles) including a 20% PhiX spike in (PhiX Control v3 Illumina).

### Long- and short-read metagenomic sequencing

Four additional stool samples from The Mucosodom human study participants, under approved ethics (REC: BAC.004.A2, 2020/21-006) were preserved with (S) and DNA extracted with Protocol 3 i.e., bead-beating via Lysis Matrix E (MP Biomedicals) and Vortex adapter for 24x2ml tubes for Vortex-Genie™2 (10 min, highest speed) (Scientific Industries SI™) and the Maxwell PureFood GMO and Authentication Kit (Promega, AS1600), and sequenced using three technologies:

1. Genomic DNA was normalised to 5ng/µl with EB (10mM Tris-HCl). 0.5 µl of Tagmentation Buffer (TB1) was mixed with 0.5 µl Bead Linked Transposomes (BLT) (Illumina) and 4 µl PCR grade water in a master mix and 5ul added to a 96 well plate. 2 µl of normalised DNA (10ng total) was pipette mixed with the 5 µl of the tagmentation mix and heated to 55 ⁰C for 15 minutes in a PCR block. A PCR master mix was made up using 10 µl KAPA 2G Fast Hot Start Ready Mix (Merck) and 2 µl PCR grade water per sample. 12 µl of this mastermix was added to each well to be used in a 96-well plate. 1 µl of 10µM primer mix containing both P7 and P5 Illumina IDT UDI (Unique Dual Index) 10mer indexed primers were added to each well. Finally, the 7 µl of Tagmentation mix was added and mixed. The PCR was run with 72°C for 3 minutes, 95°C for 1 minute, 14 cycles of 95°C for 10s, 55°C for 20s and 72°C for 3 minutes. Libraries were pooled following quantification in equal quantities. The final pool was double-SPRI size selected between 0.5 and 0.7X bead volumes using sample purification beads (Illumina® DNA Prep, (M) Tagmentation (96 Samples, IPB)). The final pool was quantified to calculate the final library pool molarity and run with final concentration of 750 pM on an Illumina Nextseq20000 instrument.
2. PacBio shotgun libraries for long-read sequencing were prepared using the SMRTbell prep kit 3.0 (PacBio, 102-182-700) and the barcoded overhang adapter kit 8A (PacBio, 101-628-400), using manufacturer’s instruction for constructing whole genome sequencing (WGS) libraries from metagenomic DNA; DNA was sheared by using Covaris g-TUBEs (Covaris, LLC, 520079) for two times at 11000 rpm for 30s and ligation was extended by incubating samples at 4°C for ∼16 hours. Additionally, as per manufacturer suggestion, AMPure PB (PacBio, 100-265-90) bead size selection was performed for the final step of library construction (size selection > 5kb). 3) ONT shotgun library preparation was carried out using the ligation sequencing kit SQK-LSK114 (Oxford Nanopore Technologies), as per manufacturer’s protocol, without size selection or shearing, and with the Native Barcoding Expansion 96 kit (EXP-NBD196, Oxford Nanopore Technologies). Libraries were sequenced on a PromethION instrument using R10.4 flow cell; raw sequencing data was collected with ONT MinKNOW software (v4.0.5). Subsequently, base calling and de-multiplexing was carried out on the instrument using guppy_build_version 7.3.1+f2bd36c15; basecall_model_version_id=dna_r10.4.1_e8.2_400bps_sup@v4.3.0.

### Bioinformatics: Read processing, assembly and binning

The 16S rRNA sequences were processed using LotuS2 v2.34.1 [18], sequences clustered into amplicon sequence variants (ASVs) using DADA2 v1.26.0 [19] analysis and a taxonomy assigned using KSGP [20] , rarefactions were conducted using rtk2 v2.11.2 [21].

All metagenomic processing from raw reads to metagenomic species (MGS) abundance was conducted using the MG-TK pipeline (https://github.com/hildebra/mg-tk). Briefly, raw shotgun metagenomes were quality filtered using sdm v3.08 [22], with default parameters, and Kraken2 [23] was used to remove human reads. Host-filtered metagenome reads, Illumina short-read data were assembled using MEGAHIT v1.2.9 [24] with parameters “--k-list 25,43,67,87,101,127”, and for ONT and PB long-read sequences, we used metaMDBG v1.1 [25] with default parameters for assembly. Reads were mapped onto assemblies using Bowtie2 v2.5.4 [26] with parameters “-- end-to-end “, genes predicted with prodigal v1.0.1 [27] with parameters “-p meta” and a gene catalogue clustered at 95% nt identity using MMseqs2 v16-747c6 [28]. MAGs (metagenome assembled genomes) were binned using SemiBin2 v1.5.1 [29] and combined in MG-TK to MGS, relying on canopy clustering (https://github.com/hildebra/canopy2) [3]. MAGs completeness and contamination were estimated using checkM2 v1.0.2 [30]. Matrix operations were carried out using rtk2 v2.11.2 [21]. Abundances of MGS in different samples were estimated based on conserved marker genes and their median abundances within the gene catalogue.

### Data analysis

Statistical analysis and visualization were done in R version 4.3.3 (R Foundation for Statistical Computing, Vienna, Austria) [31]. Alpha-diversity and Beta-diversity indices and compositional analyses were calculated using the R-packages rtk [21] and phyloseq [32] and Vegan [33].

For Alpha-diversity measures (i.e., indices of diversity), richness (observed), evenness (Shannon) and Shannon diversity index, sample count matrices were firstly rarefied to the minimal sample sum to ensure even sampling depths using rtk package. The normality of the alpha diversity distribution at different taxonomic levels was tested using a Shapiro–Wilk test implemented in the “shapiro.test” function in stats package in R. To test the statistical significance of alpha diversity across variables (Preservation methods and Protocols by different stool samples) for normally distributed data, we used Analysis of variance (ANOVA) implemented in R (“aov” function in stats package) and pairwise comparisons with Tukey’s HDS (“TukeyHSD” function in stats package). We fitted a generalized linear model implemented in R (“glm” function in stats package) with a gaussian distribution to model the alpha diversity variation between Preservation methods and Protocols by different stool samples. We used the Visreg package [34] in R to visualize the generalized linear model. The statistical significance of the models was tested using an ANOVA implemented in R (“anova.glm” function in the stats package).

For Beta-diversity (i.e., the quantification of sample dissimilarity), Bray–Curtis dissimilarities as well as taxonomic composition related analysis, taxa count matrices were normalized by dividing each feature by the respective total sample sum (TSS). To identify drivers of beta diversity, we tested whether the dispersion among groups was homogeneous using the function “betadisper” implemented in the Vegan package. Later, we assessed the statistical significance of the beta diversity among Preservation methods and Protocols by different stool samples using a Permutational Multivariate Analysis Of Variance (PERMANOVA) implemented in the “adonis2” function in Vegan package, using 9999 permutations. To investigate which metadata covariates contribute to the variation in the microbiota community, we used conditioned distance-based redundancy analysis (dbRDA, function “dbrda” in Vegan), based on Bray–Curtis distance, using the “dbrda” function in the vegan R package. Covariates found to significantly contribute to the ordination outcome were further implemented with both direction model selection on dbRDA using the “ordiR2step” function in the vegan package to determine the nonredundant cumulative contribution of metadata variables to the variation (stepwise dbRDA), as described previously [35, 36].

For univariate tests, features from the abundance matrix were removed that were present in less than four samples or had less than 0.001 relative abundance. To account for the “sample difference-effect”, significance between different features was tested with “independance_test” function in coin package [37] with teststat = “quad“ for Wilcoxon rank-sum test or “scalar” for Kruskal-Wallis test, with block for sample difference-effect, followed by a multiple testing correction (Benjamini–Hochberg). Data were visualized with ggplot2 [38] and custom R scripts.

## Results

### DNA Extraction, Sample Storage, and Their Interaction Affect the Recovery and Integrity of DNA

#### Experimental design overview

We benchmarked three DNA extraction kits, with eight different Protocols, in combination with five faecal sample storage methods, totalling to N=32 combinations, each tested in duplicate (Methods, Fig 1, Suppl. Tables 2 and 3). Three commercially available DNA extractions were used in combination with different Protocols: Maxwell PureFood GMO and Authentication kit (for brevity “MPureFood” from hereon, with Protocols 1, 2 and 3), Maxwell RSC Faecal Microbiome ( “MFaecal”, with Protocols 4, 5, 6, 7), and the QIAmp PowerFaceal pro kit (“QIAFeceal”, with Protocol 8). We refer to these Protocols throughout the text as “_1” to “_8”. Two technical replicates are denoted as “_A”, “_B”. To benchmark these Protocols, we used two stool samples from two individuals (“V1” and “V2”) corresponding to Bristol stool chart of Type 2 (“BSS 2”) and Type 4 (“BSS 4”), respectively. Five stool preservation methods were tested in combination: 1) immediately processed samples (“I”), 2) frozen samples (“FR”), 3) anaerobic storage at RT and 30°C using GutAlive Anaerobic Microbiome Collection Kit (“G”), and 4) nucleic acid preservations at RT and 30°C using OMNIgeneGUT Stabilization Solution (“O”) and 5) DNA/RNA Shield Stabilization Solution (“S”). For example, “V2FR_4_A” refers to DNA extracted from second participants (V2) stool sample, which was frozen (FR) prior DNA extraction using MFaecal kit and Protocol 4 (_4), with (_A) indicating the first technical replicate.

**Fig. 1.**
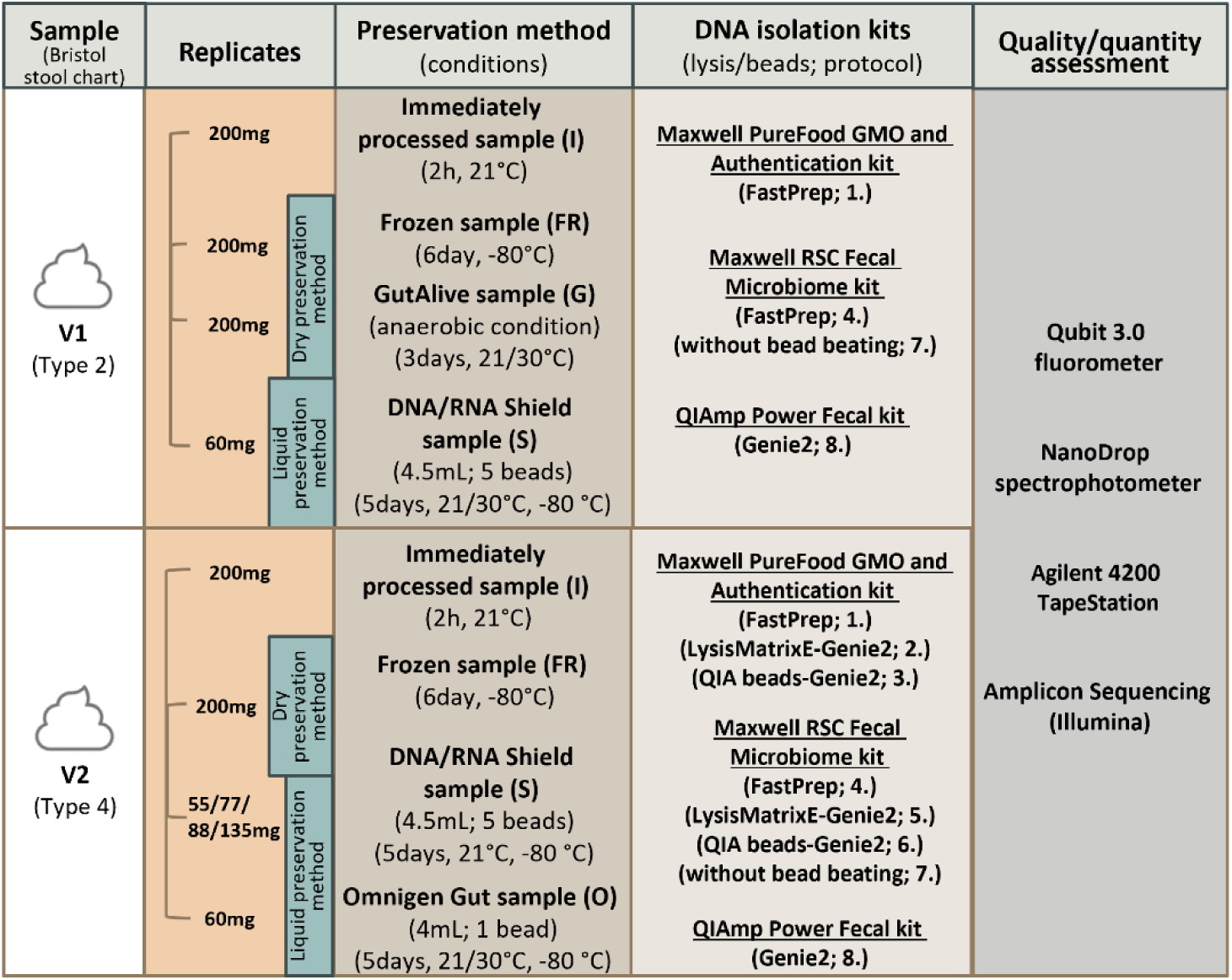
Benchmarking of protocols. We used two different human stool samples that were tested in a combination of 32 different approaches. Stool V1 was Bristol stool chart type 2, Stool V2 was type 4, representing a spectrum of stool consistencies. A combination of preservation methods (column 3) and DNA isolation kits, including different cell lysis methods (column 4) was tested for each and evaluated using different approaches (column 5).

### DNA yield and quality vary between extraction kits and different preservation approaches

To ascertain the suitability of different protocols and extraction methods for long-read sequencing, we first analysed the quantity and quality of DNA isolated from each extraction protocol using both fluorometric (Qubit) and spectrophotometric (NanoDrop) methods (Fig.2). There was no statistically significant difference between quantity of the extracted DNA using different protocols, preservation or extraction kits (Kruskal-Wallis, BH adjusted p-value: >0.05: Fig.2 A). The highest yield for the first stool (V1) was obtained by using protocol 1, preserved with (G) (median 70.30 ng ± 18.9), and for the second stool (V2) was protocol 8, preserved with (O) (median 106.80 ng ± 5.2). For both stool samples (V1 and V2), the lowest DNA yield came from enzymatic lysis (Protocol 7, median 4.65 ng ± 0.97; Suppl. Table 4). In general, protocols using mechanical lysis yielded significantly higher DNA concentrations, regardless of the bead-beating type and composition of the beads (Signed Rank Test, BH adjusted: enzymatic vs mechanical/chemical lysis p-value: 0.0002 and enzymatic vs mechanical/enzymatic lysis p- value: 0.055). All the extraction methods yielded sufficient DNA concentrations for either ONT or PacBio sequencing, except for Protocol 7 (median 4.65 ng ± 0.97) and the combination of (S) preservation with Protocol 8 (median, 1.10 ng ± 0.28).

**Fig. 2.**
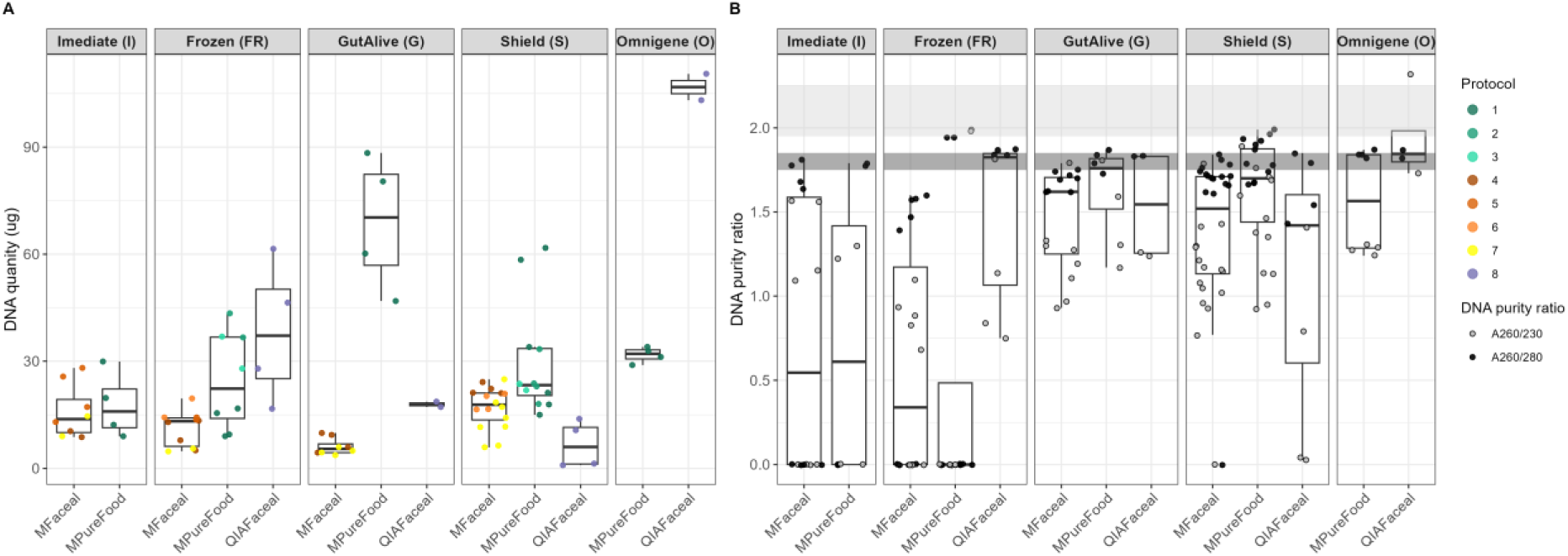
(A) Comparison of DNA concentration/quantity and (B) purity at OD 260/280 and OD 260/230 between different preservation method and extraction protocols (dark grey shaded area represent the recommended A260/280 ratio and light grey area represent the recommended A260/230 ratio).

To assess DNA purity and identify potential non-nucleic acid contaminations, we analysed absorption spectra with NanoDrop Spectrophotometer (Fig.2 B, Suppl. Table 5). Most DNA extracts were below the optimal A260/230 and A260/280 ratios (2.0, 1.8, respectively). The highest A260/230 ratio was achieved with protocol 1 (median: 1.28 ± 0.6), and highest A260/280 ratio with protocols 2 and 8 (median: 1.83 ± 0.98 and 1.84 ± 0.14, respectively). In general, stool preservatives showed a better A260/280 and A260/230 ratio than frozen samples (Signed Rank Test, BH adjusted p-value: A260/280: (O) vs (FR) 0.046; (S) vs (FR) 0.012; A260/230: (O) vs (FR) 0.004 and (S) vs (FR) 0.0005). However, all of the DNA extracts fall below the official purity recommendation for ONT and PacBio library construction (*20*).

Next, we investigated DNA fragment size distribution among protocols and preservation methods, using Tapestation electrophoresis (Fig.3). Firstly, we assessed the DNA integrity (DIN) value, a measure for determining the integrity (e.g, degradation and shearing) of gDNA (Fig.3 B, Suppl. Table 6). Statistically significant differences in DIN values were observed between protocols, preservation or lysis method (Kruskal-Wallis, BH adjusted p-value: 0.047, 0.008 and 0.043, respectively). Protocols using only enzymatic lysis yielded in significantly higher DIN values (Protocol 7, median 8.35±0.9) compared to mechanical lysis (Signed Rank Test, BH adjusted: enzymatic vs mechanical/chemical lysis p-value: 0.016; Fig.3 B). Furthermore, DNA isolated from samples preserved via (S) and (O) retained the highest DNA integrity (median 9.5 ± 0.75 and 8.10 ± 0.09, respectively) in comparison to (FR) storage conditions, which had the highest degree of shearing leading to diminished DNA integrity (median 6.60±0.4).

**Fig. 3.**
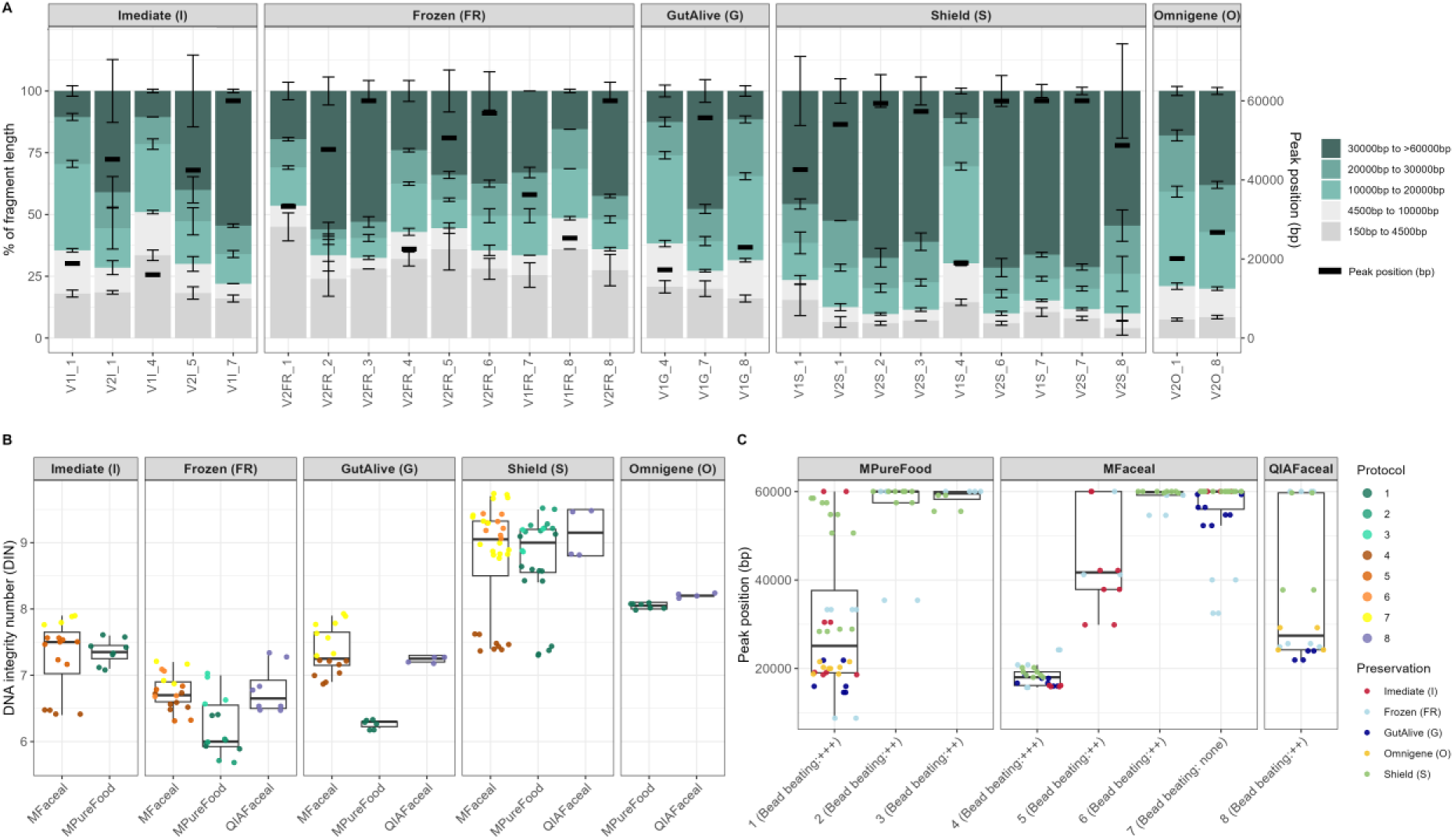
(A) Fragmentation profiles of DNA extracted by different isolation kits. The different user defined size classes are plotted as stacked bars representing mean % of total DNA between replicates. Error bars represent SD between replicates. (B) DNA integrity values (DIN), representing the amount of degradation and shearing of the DNA extracts between different preservation methods and DNA extraction kits. Colour of dots represent the different protocols used. (C) Peaks positions of fragmentation profiles (the centre of the regions’ mass length in bp) between different DNA extraction kits and extracted by different protocols. Colour of dots represent the different preservation method used, and the comments beside the protocol number on x axis represent the amount of bead beating used in each protocol (“+++” - FastPrep; “++” - Genie2; “none” - no beating used). Samples with DNA concentrations <0.4 ng μL^−1^ were excluded from the analysis.

Secondly, we assessed the Tapestation peaks, which represent the highest concentration of extracted DNA at given size (Fig.3 A). In general, there was no statistical difference between peaks of fragment lengths of the extracted DNA using different protocols, preservation or extraction kits (Kruskal-Wallis, BH adjusted p-value: >0.05). However, as previously observed with DIN values, protocols using enzymatic lysis yielded longer DNA, compared to samples processed with mechanical lysis (Signed Rank Test, BH adjusted p-value: enzymatic vs mechanical/chemical lysis: 0.008; and enzymatic vs mechanical/enzymatic lysis: 0.001). For example, comparing the peaks between Protocol 1, 4 or 8 using mechanical shearing for longer times, we obtained peaks at statistically shorter fragment lengths than by using Protocol 7 (Signed Rank Test: p-value 0.0004, 0.0001, and 0.014 respectively; Fig.3 C). Therefore, the longest fragment length peaks were obtained with Protocol 7 (median peak 60,000±8145 bp; Fig.3 C). It is noteworthy that (S) preserved DNA had consistently larger fragment sizes with 64.5% of extracted DNA >30k bp (median peak 59.7kbp ± 15.3kbp; Fig.3 A) regardless of the extraction protocol used (Kruskal Wallis, BH adjusted p-value: >0.1; Fig.3 B).

The identified combination (Protocol 3 with DNA/RNA Shield preservation) was applied to single isolates of Gram –, Gram + and a mock community of bacteria (2x Gram +, 4x Gram –), to validate the generalized applicability (Suppl. Table 13, Suppl. Fig.4). This showed similar results, with high DNA concentrations extracted (Gram –: 156 ± 29.60 ng/µL, Gram +: 262 ± 83.76 ng/µL , mock: 238 ± 53.33 ng/µL) and high DNA integrity numbers (Gram –: 9.3 ± 0.44 ng/µL, Gram +: 9.2 ± 0.26 ng/µL, mock: 9.2 ± 0.17 ng/µL). This was reflected in long DNA fragments being extracted, with 50.6 ± 5.75% (Gram –), 55.90 ± 8.74% (Gram +), and 54.57 ± 4.84% (mock) of total DNA above 30 kb in length. The median peak fragment sizes were similarly high (Gram –: 51,375 ± 4,211 bp, Gram +: 50,807 ± 4,729 bp , mock: 50,211 ± 2,360 bp), validating the favourable results obtained in our faecal DNA extractions.

**Fig. 4.**
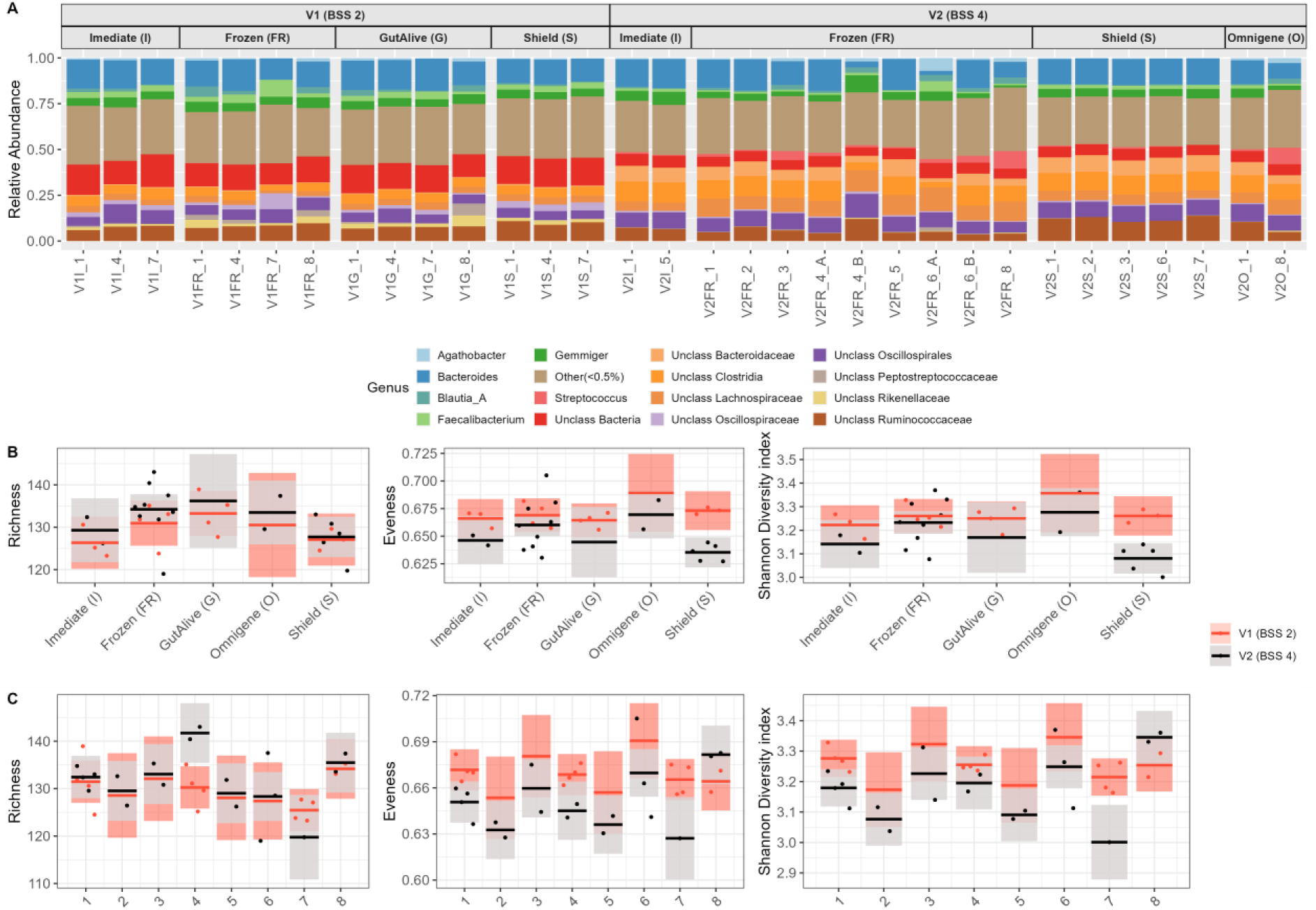
Effect of the preservation method and extraction protocols on the taxonomic profiles of the samples on genus level. (A) The taxonomic classification of the main taxa (taxa with relative abundance below 0.05 were grouped and named “Others”) identified in the faecal samples preserved with different preservation methods and DNA extraction protocols. (B, C) Genus-level alpha diversity (Richness, Evenness and Shannon diversity index) compared among different (B) preservation methods, and (C) different protocols; colours represent different participants stool sample, smooth areas represent the confidence bands, dots represent the partial residuals, and lines the predicted values inferred from the models.

Current recommendation for PacBio and ONT sequencing is at least 1µg of average >30 kb of total DNA with 260/280 nm ratios between 1.8-2 and 260/230 nm ratios between 2.0-2.2 for ONT and 1.25 - 4µg (20-50ng/µl in 25µl or 40-80ng/µl in 50µl) of 70% of DNA ≥10 kb for PacBio. Based on the substantial DNA yield, quality, uniform length distribution and ease of sample collection, our data suggests that protocol 3 with DNA/RNA shield (S) preservation is most suitable for long-read metagenomics.

### Taxonomic consistency of different sample treatments

Initially, we evaluated the taxonomic consistency between technical replicates across all protocols and preservation methods and assessed the impact of stool preservation methods on taxonomic consistency for samples, which vary in storage temperature and stool input quantity (i.e., samples preserved with (S), (G) and (O) at RT and 30°C, and samples preserved with (S) for stool input quantity). No significant differences were observed between technical replicates, except for V2, preserved at -80 °C and extracted by using protocols 4 and 6 (Wilcoxon sum rank: p-value > 0.05; Suppl. Fig.1). Regarding the samples stored at different temperatures, no significant difference was observed between samples stored at RT or 30 °C, except in sample V1 preserved with (G) and extracted by using either protocols 1 or 4 (Wilcoxon sum rank: p-value < 0.05; Suppl. Fig.2). No significant difference was observed between different stool quantity as an input for DNA extractions (Kruskal Wallis p-value: > 0.05; Suppl. Fig.3). Based on these results, in the following analysis the median values of all technical replicates are used. we used the median value for all different stool concentration, samples stored at different temperatures and technical replicates for further figure creations and readability.

To investigate the effect of preservation and DNA extraction on taxonomic representation (Fig.4), all samples were sequenced via 16S amplicon sequencing. To model the variation of alpha diversity across different preservation methods or protocols and two different sample types i.e, V1 and V2, we constructed Generalised linear models (GLMs) using a gaussian distribution based on normal distribution of Richness, Evenness and Shannon diversity index at genus level (Shapiro–Wilk, p-value > 0.05). Using this approach, we found significant differences in the alpha diversity of microbial communities between different sample types and preservation methods (ANOVA: eveness p = 0.004, F = 10.13; and Shannon diversity index p = 0.008, F= 8.29), however only Shannon diversity differed significantly between preservation methods (S) and (Fr) in sample V2 (TukeyHSD, p-value < 0.05; Fig.4 B). Regarding the differences between protocols, alpha diversity (richness) was different (ANOVA: p<0.05) between protocols 4 and 7, and for Shannon diversity index between protocols 7 and 1, protocols 8 and 7, and protocols 8 and 5 (TukeyHSD, p-value < 0.05; Fig.4 C and Suppl. Table 7).

Next, we investigated how much community composition was affected by the tested protocols, visualizing differences in a principal coordinate analysis (PCoA). This showed that samples clustered primarily by the donor of faecal samples (PERMANOVA p <= 0.001, F=114.3; Fig.5 A). When accounting for differences between the stool participants (V1 and V2), the microbiome composition was significantly different between different extraction protocols (PERMANOVA p <= 0.001; F = 5.95) and preservation methods (PERMANOVA p<= 0.001; F = 8.06; Suppl. Table 8). The homogeneity of between group dispersions (betadisper) was significantly different between preservation methods (ANOVA: V1 p = 0.0019, F = 6.11; and V2 p = 0.0012, F= 6.49), with (S) having the smallest (0.044 for V1, 0.033 for V2), and (FR) having the largest dispersion (0.132 for V1 and 0.141 for V2; Suppl. Table 8), meaning the variability in taxonomic composition were highest among used protocols, when relying on (FR) samples. Comparing extraction protocol and preservation method in relation to the baseline condition (i.e. Immediately (I) extracted sample, with protocol 1), (FR), as well as samples extracted using protocol 8, had a higher distance to baseline (Fig.5 B), further confirming our observations.

**Fig. 5.**
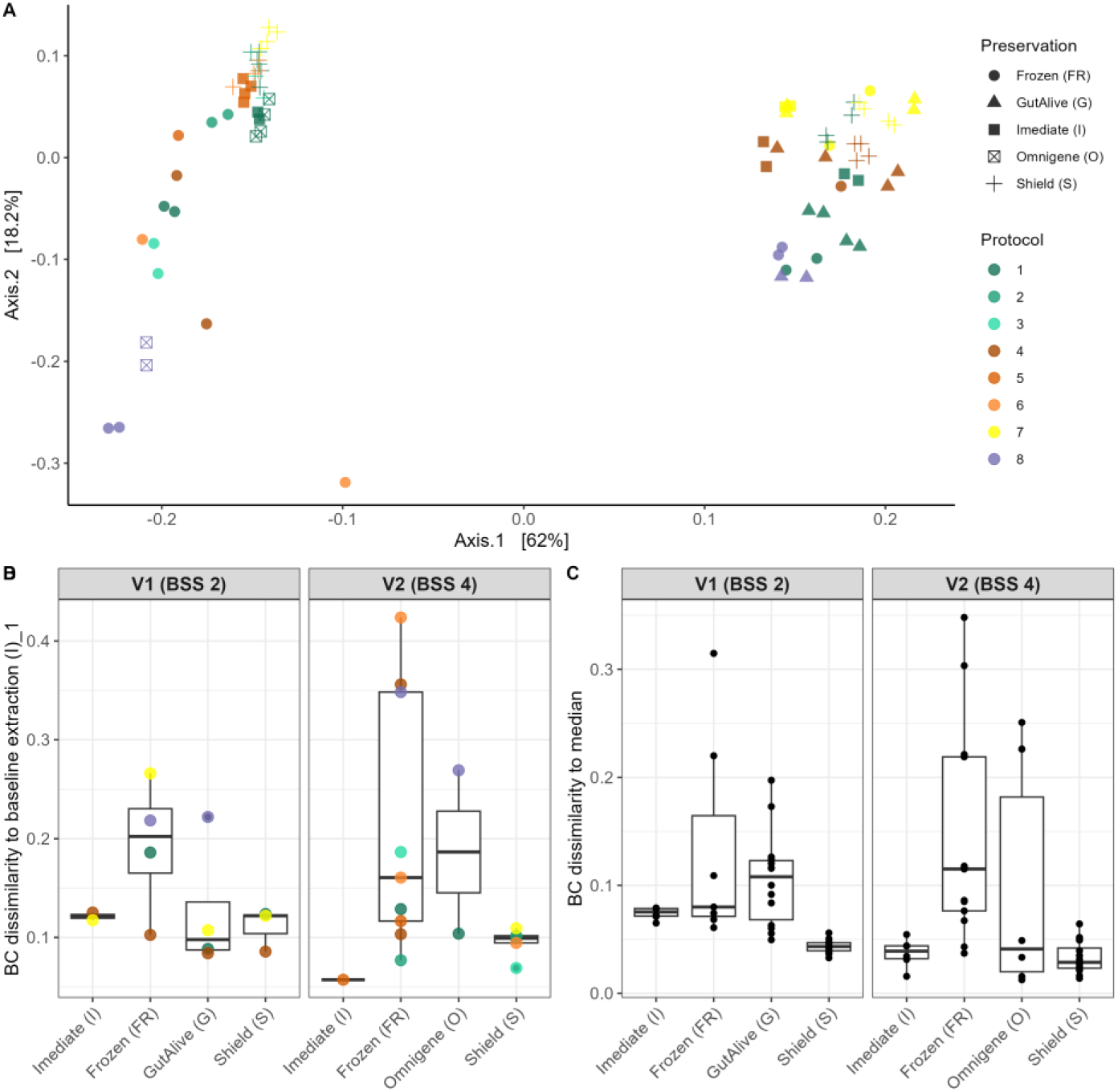
Multivariate analysis at genus level. (A) A principal coordinate analysis (PCoA) using Bray-Curtis dissimilarities of the faecal samples with different preservation methods and extraction protocols. (B) Bray-Curtis dissimilarities to the baseline condition ([(I)_1]; i.e., immediately extracted sample with protocol 1) and other protocols. (C) Bray-Curtis dissimilarities to the median among samples extracted with different protocols but the same sample preservations in order to describe homogeneity of group dispersions.

The relationship between experimental conditions and genus-level microbiome composition was visualized using distance-based redundancy analysis (dbRDA; Fig.6). This approach enabled us to estimate both the independent contribution (i.e., individual explanatory power) and their cumulative contribution (i.e., cumulative explanatory power) of each variable. Independently, interindividual variation accounted for the largest portion of explained variance (R² = 60.2%). Among the other experimental variables, protocol contributed R² = 30.7% and preservation method R² = 26.9%, followed by lysis type R² = 12.8% and DNA extraction kit R² = 8.7% (Fig. 6C). However, analysis revealed collinearity and larger within group variability (Suppl. Table 10), particularly for variables Protocol and DNA extraction kit, as the protocol variable in our study inherently combines both DNA extraction kit and lysis type.

**Fig. 6.**
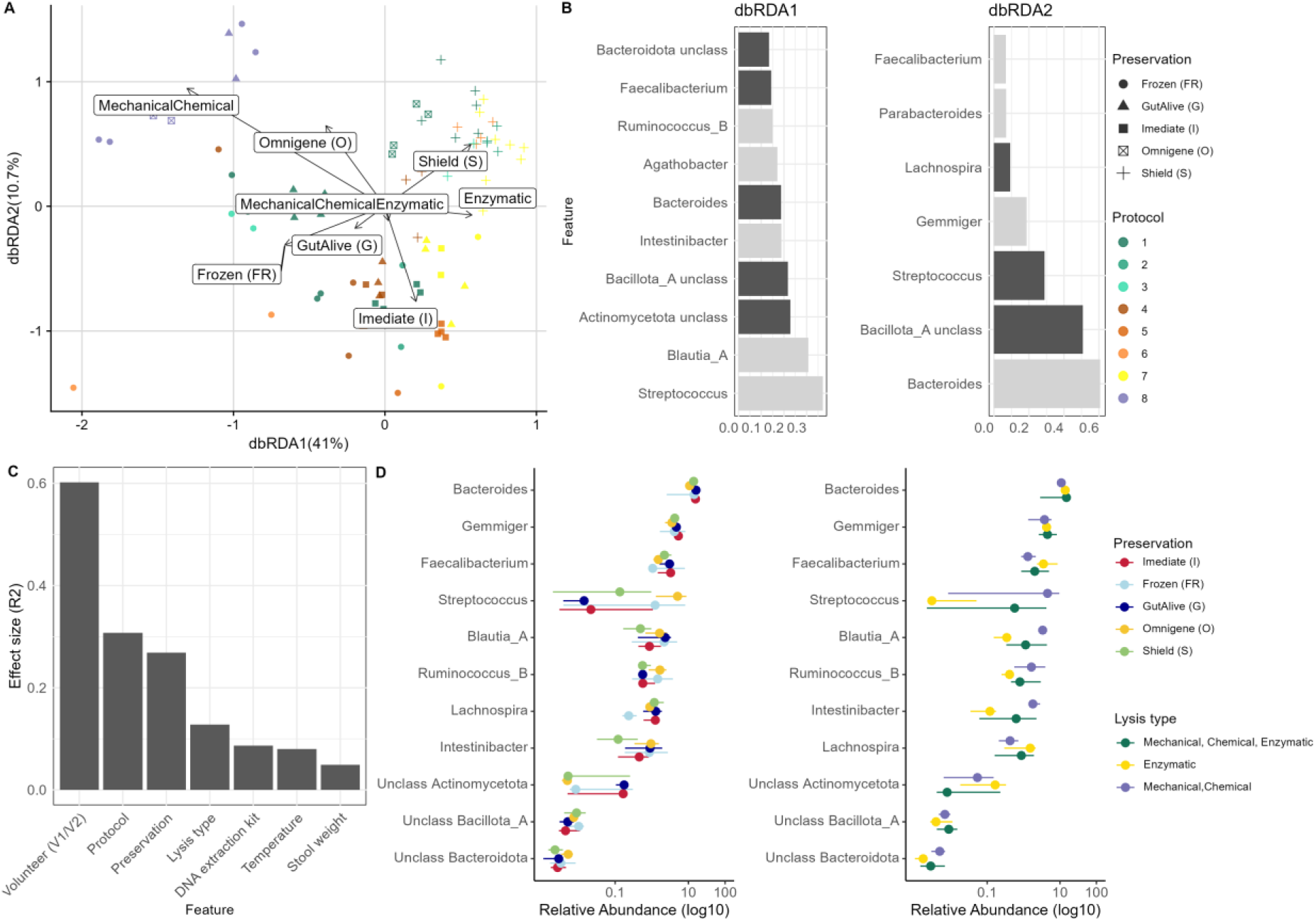
Taxa correlating to Preservation and Lysis type on genus level. (A) Constrained ordination (distance-based redundancy analysis [dbRDA]) using Bray-Curtis dissimilarity and conditioned for participants, with arrows representing different Preservation methods and Lysis type and their relation to sample composition. (B) Visualization of the most strongly correlated taxa to dbRDA axis 1 and 2. (C) The individual explanatory power of the variables (dbRDA R^2^). (D) Relative abundance of correlated taxa identified by dbRDA among Preservation methods (left graph) or Lysis type (right graph); dots represent the median and lines the upper and lower quantiles, colours representing either different Preservation methods or different Lysis type. All taxa were significantly different between groups of methods (Kruskal-Wallis, blocked by interindividual variation, BH adjusted p-value < 0.05).

To reduce redundancy and improve interpretability, the final model focused on preservation method and lysis type, which together explained R² = 18.4% of the variation in microbiome composition, while conditioning for interindividual variation (Suppl. Table 10). Axis 1 of the dbRDA correlated mostly with mechanical/chemical vs enzymatic lysis, while axis 2 correlated with samples that were immediately (I) processed vs preserved with (O) and (S) (Fig.6 A).

The axis 2 on dbRDA (Fig.6 A) revealed distinct groups of samples that were treated with only enzymatic lysis (positive effect direction) or a combination of Mechanical/Chemical lysis with protocol 8 (negative effect direction). Axis 1 of the dbRDA correlated to liquid preservation and anti-correlated with dry preservation methods. Interestingly, samples preserved with (O) when extracted with protocol 8 had a negative effect direction on axis 2, but when extracted with protocol 1 had positive direction. Similar trends can be observed on axis 1 for (FR) or (G) preserved samples.

The identified taxa correlating most strongly to the ordination were Bacteroides, Gemmiger, Streptococcus, Ruminococcus_B, Lachnospira, Intestinibacter, Faecalibacterium, Blautia_A, Agathobacter, and unclassified Bacteroidota, Bacillota_A, and Actinomycetota, which belong to the Phyla Firmicutes, Bacteroidota, Bacillota, and Pseudomonata. Of note, all taxa correlated to the dbRDA are Gram positive bacteria and known to be difficult to lyse and extract the DNA (Fig.6, B and D), except for Gemmiger and Bacteroides.

Similar taxa were also identified using univariate testing for differences among preservation methods and lysis type, e.g. Phyla Bacillota_A was significantly different between (FR), (G) and (S), and between different lysis type (Suppl. Table 11). Furthermore, genera Phocaeicola and Copromonas differed in relative abundance between all preservation methods, however only Clostridium, Enterocloster, and Anaerostipes were different between liquid preservation methods (O and S) and dry preservation methods (Suppl. Table 8). Interstingly, genus Alistipes, Intestinimonas, Dysosmobacter and Fecalibacterium were more prevalent in samples were only enzymatic lysis was used.

Based on the stable taxonomic profiles obtained and the previously reported DNA purity and length characteristics, we can further confirm protocol 3 with DNA/RNA Shield (S) preservation is most suitable for long-read metagenomics

### Protocol validation in faecal long-read metagenomics

To validate our findings, four human faecal samples were processed with protocol 3 with (S) preservation, yielding indeed high-quality DNA at high molecular weight (Fig.7 A). Subsequently, samples were sequenced using a) short-read Illumina NovaSeqX, b) long-read ONT Promethion and c) PacBio Revio. Sample collection, DNA extraction and library construction proceeded with minimal efforts, the yields per platform were 9-15 Gbp (Illumina), 3-8 Gbp (ONT), and 6-9 Gbp (PacBio) per sample, amounting to 87M (Illumina), 2M (ONT), 1M (PacBio) reads/sample. Metagenomes were processed with the MG-TK pipeline [39]. ONT metagenomes had the highest % of human reads (1.6%), Illumina intermediate (0.37%), while PacBio was lowest (0.03%) despite an otherwise equal sample treatment. After assembly, we obtained on average N=20, 21 and 32.5 high quality bins /sample for Illumina, ONT and PB, respectively. However, using metaMDBG [25] we assembled more complete circular contigs (CC) for ONT then PacBio samples (average 54.8 and 48.3, respectively), with none for Illumina as one would expect for short-read sequencing. However, most CC probably represented plasmids, as the number of CC’s ≥1Mb in size was 6 for ONT and 8.3 for PacBio (Suppl. Table 9).

**Fig. 7.**
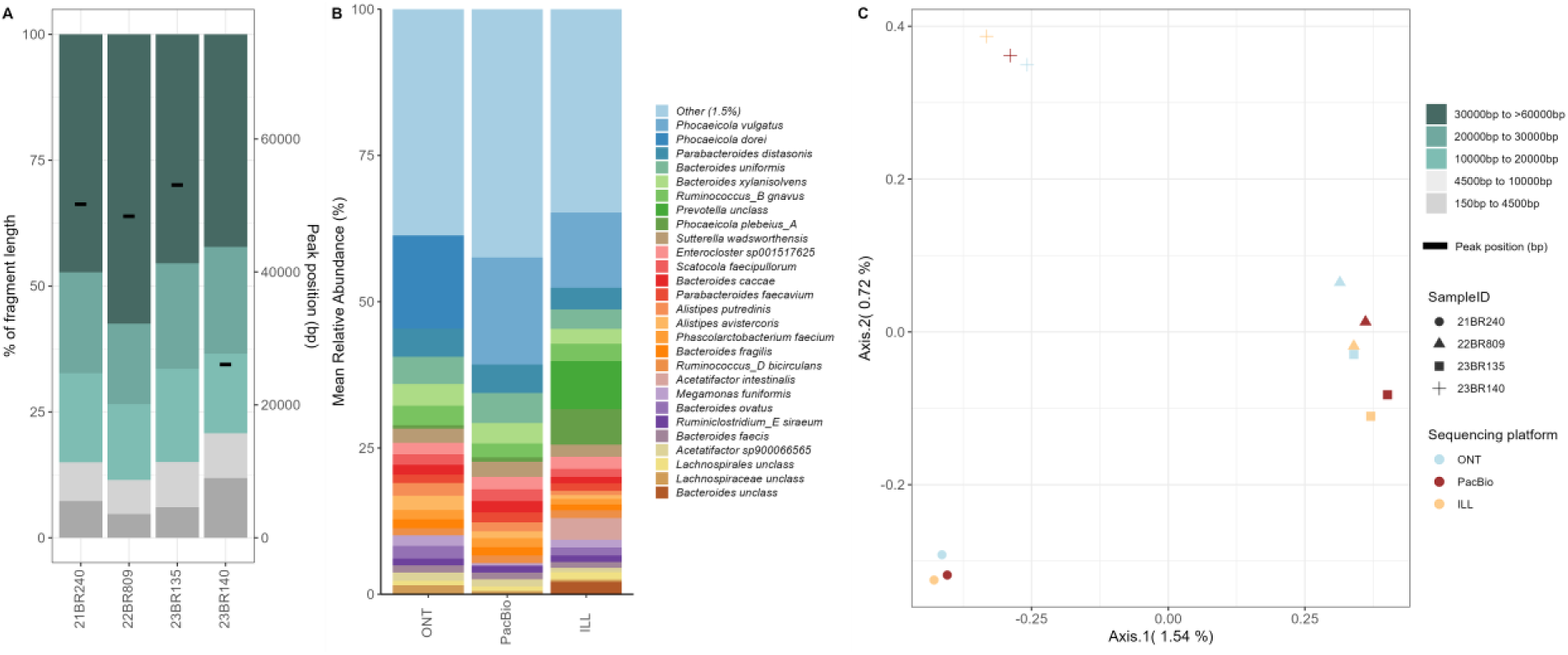
Validation of protocol 3 with DNA/RNA Shield (S) preservation on four faecal samples. (A) The fragmentation profiles of extracted DNA from validation samples. (B) Mean relative abundances (averaged across 4 samples) of dominant taxa sequenced with different platforms (taxa with relative abundance below 0.015, were grouped and named “Others”). (C) Principal-coordinate analysis (PCoA) (Bray-Curtis dissimilarity) of species level profiles from validation samples. Inter-sample differences were driving microbiome variation, with little variation introduced by the different sequencing platforms.

MG-TK [39] metagenome processing demonstrated the reconstruction of complete genomes, while the overall community composition was mostly conserved between sequencing platforms (Fig.7 B). Most of the variation in community composition was driven by inter-sample differences (PERMANOVA p=0.001, F=13.2), but choice of sequencing platforms also introduced significant differences in overall composition (PERMANOVA p-value= 0.004; F=0.68; Fig.7 C). In total more species were detected with PacBio sequencing in comparison to ONT and Illumina (n=344, n=285 and n=312, respectively), only *Phocaeicola dorei* and *Phocaeicola vulgatus* being significantly different in relative abundance between platforms (Wilcoxon sum rank: p-value: 0.02 and 0.03, respectively); however, the relatively small sample size could have masked further variation. Nonetheless, the overall stable community profiles between technical replicates, clear distinction between biological samples, large DNA molecules and efficient recovery of almost complete genomes from metagenomes demonstrated the quality and applicability of the here presented protocol combination, suitable for small- and large-scale sequencing projects.

## Discussion

Accurate measurements of microbial communities are key to understanding bacterial communities, and DNA sequencing-based approaches are currently the most important tool for this task. Despite high-throughput sequencing (amplicon and shotgun Illumina sequencing) providing considerable insights into the human gut microbiota, many factors including sample storage prior to DNA extraction, are known to impact on the accuracy of the results achieved [10]. Characterizing and understanding such biases based on sample treatments is crucial for the design, choice of sequencing, analysis and interpretation of microbiota studies. Our work had the aim to analyse different methods for stool preservation and DNA isolation, to find the optimal combination for long-read metagenomic sequencing and unbiased microbial reporting. This showed that DNA/RNA Shield (S) sample preservation and protocol 3 (Maxwell PureFood GMO and Authentication kit, using bead-beating as the mean of mechanical lysis) were the most suitable for long-read sequencing in terms of DNA yield, quality, fragments length and taxonomic stability.

Sample preservation is essential in setting up large studies where samples need to be stored over prolonged periods to ensure standardized processing. Freezing faecal samples had been established as the gold-standard in the field, but is increasingly criticized for requiring costly logistics (“cold-chains”) [40, 41], and the storage at low temperature was reported to induce biases, such as changing the ratios of Firmicutes to Bacteroidetes [42], Lachnospiraceae to Bacteroidaceae [43], or decrease the relative abundance of Proteobacteria [16]. Significant changes in the abundance of taxa were also observed with incremental increases in freeze–thaw cycles [44]. Furthermore, our study indicates that freezing samples without including preservation liquid increases DNA fragmentation, as the quality of the DNA and DIN values were amongst the least favourable observed among all methods, concordant with previous studies [45], making freezing samples unsuitable for long-read metagenomics. The useability of frozen faecal samples without including preservation liquid to study the gut microbiome is further questioned by our investigation of microbiome reproducibility: frozen samples had a notably increased variation between technical replicates (Fig.5). And although our samples were homogenized (after defrosting), this may be insufficient, suggesting that for future studies faecal homogenization should ideally occur before the sample is frozen (though this may be difficult for practical reasons).

Using liquid preservatives to preserve faecal samples might also introduce taxonomic biases [46]. Concurrently, we observed that the taxonomic composition of dry and liquid preservatives formed distinct clusters (Fig.6). Taxonomic composition was even more influenced by the choice of lysis methods: enzymatic lysis is known to bias the Gram – to Gram + bacterial ratio, due to these two types of cell membrane being variably accessible to enzymes used to break open cell membranes [46]. Similarly, the strength and length of physical cell lysis via bead-beating can bias taxonomic representation [9], with more intensive bead beating additionally introducing a higher degree of DNA shearing [9, 47], and therefore inadvisable for long-read sequencing. To our surprise, it seemed that DNA/RNA Shield (S) preservation counteracted DNA shearing during physical cell lysis, as we consistently obtained longer DNA fragments, regardless of the amount/intensity of bead beating. We hypothesize that chelated DNA is more resilient to physical stresses introduced during bead beating, such as foaming and oxygen stress [48].

We further observed that none of our tested DNA extractions yielded DNA of sufficient purity to meet recommendations. The A260/A230 ratio, when deviating from 2.0-2.2 can signify residual chemical contamination such as EDTA, phenol, guanidine salts (often used in column-based kits) or carbohydrate carryover [11]. The ratio of A260/A280 indicates DNA purity, with pure DNA having a ratio of 1.8, while deviations can signify contamination with proteins, phenol or other molecules absorbing at or near 280 nm [11], decreasing the effectiveness of library construction and sequencing. Therefore, it is our recommendation to additionally purify DNA prior to long-read sequencing.

Interestingly, we frequently observed in our trials that DNA extraction methods have to be matched with specific sample preservation methods, which in theory could also apply/extend to stool sample type. For example, faeces preserved with DNA/RNA Shield (S) yielded less extracted DNA when using QIAGEN PowerFaceal kit (Fig.2 B). This may be due to the chelating liquid (S) being incompatible with silica column, and/or because the longer fragments obtained with (S) were not efficiently eluted into the elution buffer and instead retained on the column, likely due to their high prevalence, as the applied protocols were optimized for shorter DNA fragments (Fig. 3). And although both choice of DNA extraction kit (and consequently protocol) and sample preservation biased the taxonomic representation in our trials, the interplay between preservation and extraction method seemed important. For example, samples preserved with OmnigeneGut (O) when extracted with QIAGEN PowerFaceal kit (protocol 8) had a negative effect direction on axis 2 (dbRDA), but when extracted with Maxwell PureFood GMO and Authentication kit (protocol 1) had positive direction. Similar trends can be observed on axis 1 for frozen (FR) or GutAlive (G) preserved samples (Fig.4).

Our findings indicate that Protocol 3 and DNA/RNA Shield (S) preservation method was the optimal choice for both cultured isolate, mock community and faecal microbiome sequencing. This was validated on four faecal samples sequenced using Illumina, ONT and PacBio platforms. Sample storage, DNA extraction and library preparation were conducted effortlessly, and we observed only minor taxonomic biases between different sequencing platforms. However, of note is that ONT sequencing increased both host reads and small circular contigs, proportionally. We hypothesise, that this could be due to the size selection in the PacBio library preparation, while for ONT and Illumina library preparations did not include size selection. Human DNA in metagenomes samples is often represented as shorter fragments [8], and could thus have been removed during PacBio library preparation protocol. Similarly, the small circular contigs could have been removed in the PacBio size selection. The increase of both human reads and small circular contigs in the ONT data could be due to preferential sequencing of shorter DNA fragments [49], highlighting the need to standardized size selection steps during metagenomic sequencing experiments.

We acknowledge the limitations of this study; in that we can only comment on the global levels of the groups enumerated. In future studies, it may be interesting to examine the combined effects of sample type, preservation method and DNA extraction kit/protocol on variation in taxonomic representation on a larger set of samples to further explore our observations of those combined effects.

## Conclusion

With long-read metagenomics becoming more commonplace, standardized protocols that enable efficient and stable (meta)genomic sequencing are urgently needed. To reach these wider goals, we propose a protocol that is practical (avoiding cold chains and using robust methods), affordable, reduces taxonomic variability in technical replicates and can significantly increase the yield of high-quality, high-molecular weight DNA.

## Acknowledgements

We would like to thank all members of the Hildebrand group for valuable input at various points in the project. We would like to thank the NRP Biorepository for collection of samples and for providing long term storage of samples from the Mucosodom study, and their invaluable contributions and advice regarding participant involvement in this clinical research. We would like to thank Human studies team, for collection of the participants samples. We would also like to acknowledge all the participants involved in the study.

## Declarations

### Ethics approval and consent to participate

The Quadram Institute of Bioscience (QIB) HNU/CRF facility collected stool samples from two QIB Participant / Donor NRP biorepository participant’s database and handled under ethics approval UEA FMH ref: 201414-06HT and NNUH R&D ref: 70-05-19. Four additional stool samples used for long-read sequencing were taken from The Mucosodom human study participants database and handled under approved ethics REC: BAC.004.A2, 2020/21-006.

### Availability of data and materials

The datasets generated and/or analysed during the current study are available in the ENA repository, https://www.ebi.ac.uk/ena/browser/search; Assession number: PRJEB90666. All data analysed during this study are included in this published article (and its supplementary information files). Code is available at: https://github.com/CeKlar/HMW-DNAProject.git.

### Author’s contributions

KC: Conceptualization, Methodology, Formal analysis, Investigation, Writing - Original Draft, Visualization, Supervision

NS: Methodology, Writing - Review & Editing

AD: Methodology, Validation, Writing - Review & Editing

DB: Methodology, Writing - Review & Editing, Supervision

MK: Formal analysis, Writing - Review & Editing

JA-J: Methodology, Writing - Review & Editing

RJ: Methodology, Writing - Review & Editing

CQ: Methodology, Writing - Review & Editing

FH: Conceptualization, Writing - Original Draft, Writing - Review & Editing, Supervision, Funding acquisition

