## Supplement Figures for "Optimizing storage, high-molecular weight DNA extraction and genome reconstructions from human faecal samples"

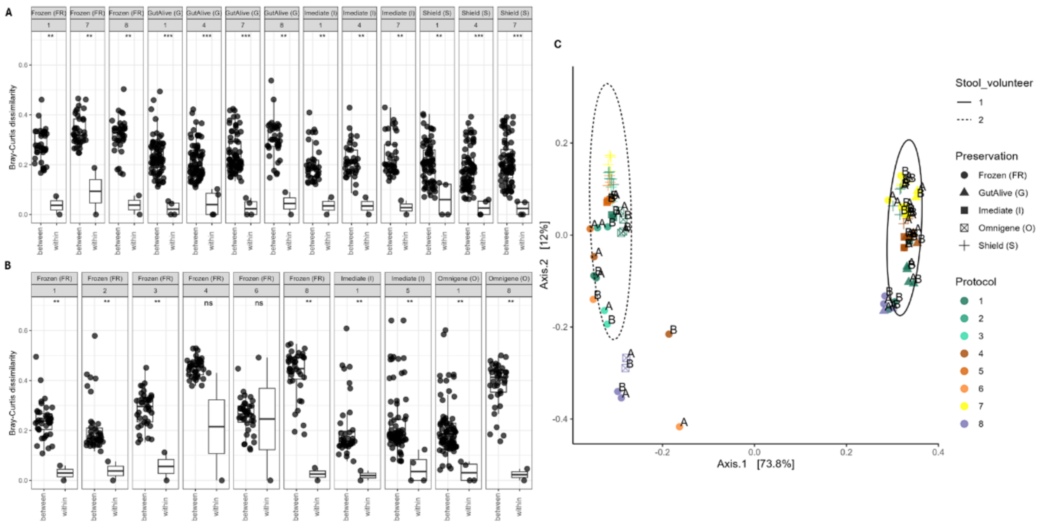


Supplement Fig. 1: Bray-Curtis dissimilarity at ASV level between and within duplicates: (A) for stool sample V1, and (B) for stool sample V2; Wilcoxon with BH adjustment (Signif. codes: 0 ‘***’ 0.001 ‘**’ 0.01 ‘*’ 0.05 ‘.’ 0.1 ‘ ns ’ 1). (C) The ordination plot (PCoA), with colours representing different protocols and shapes different preservation methods; A and B on the figure denotes each of the duplicate within different protocol and preservation used.

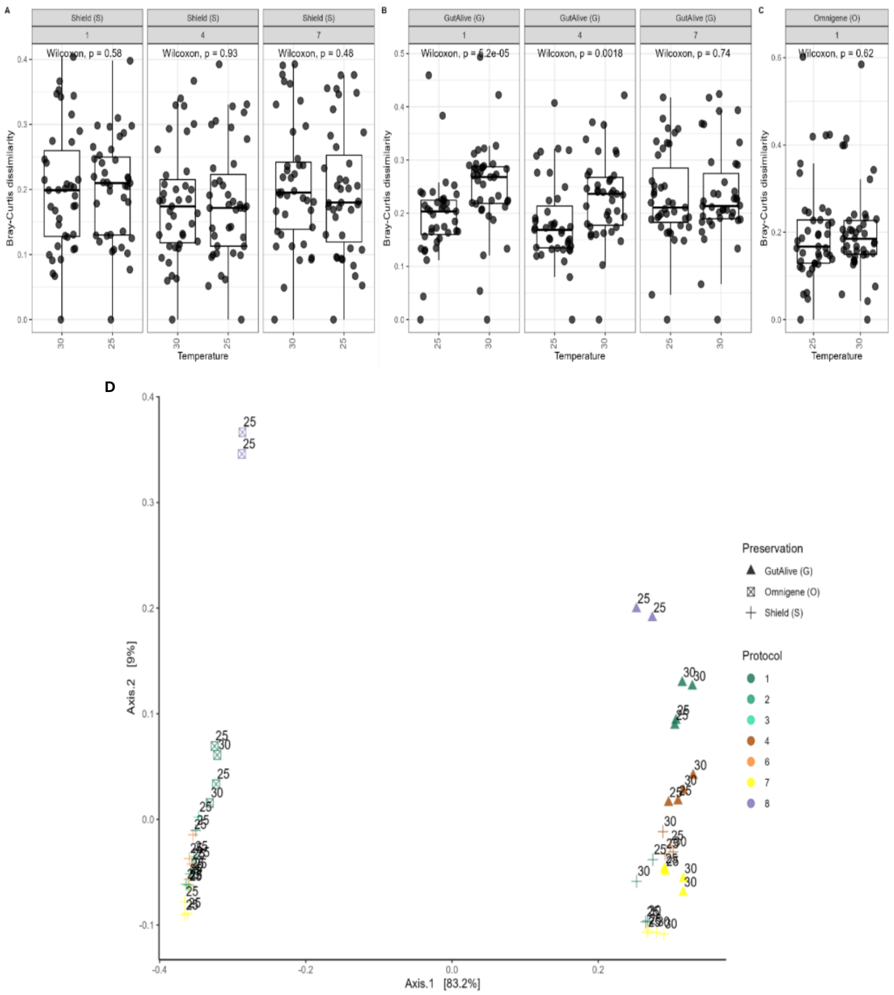


Supplement Fig. 2: Bray-Curtis dissimilarity at ASV level between different storage temperatures: (A) for preservation Shield (S), (B) for preservation GutAlive (G) and (C) for preservation Omnigene (O); Wilcoxon test with BH adjusted p-values. (D) The ordination plot (PCoA), with colours representing different protocols and shapes different preservation methods and with 25 and 30 representing each of the duplicates and temperatures.

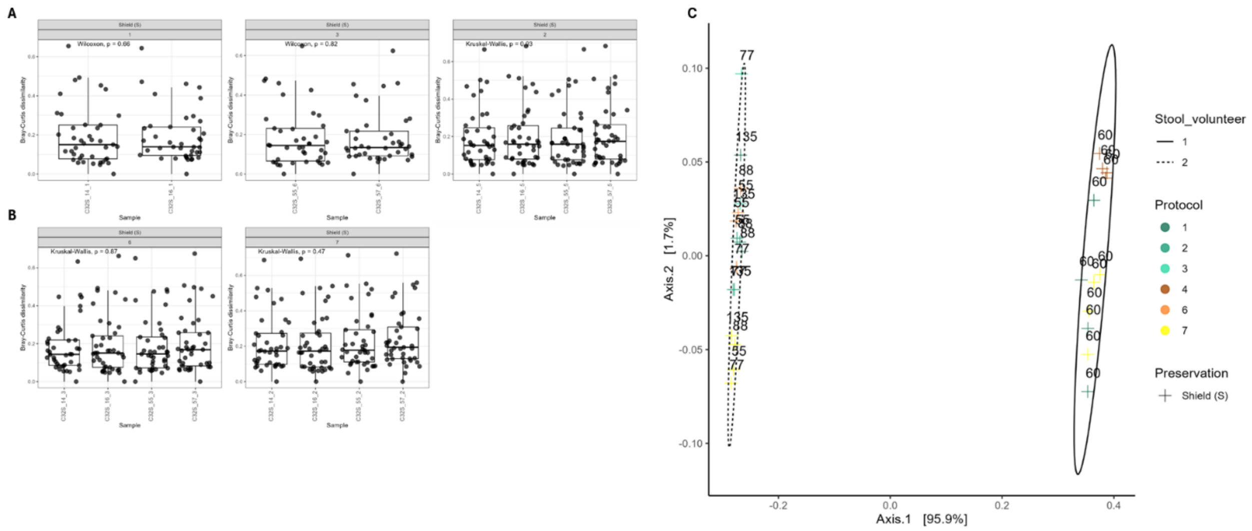

Supplement Fig. 3: Bray-Curtis dissimilarity at ASV level between stool concentration as input into DNA extraction: (A) for stool sample V1, and (B) for stool sample V2; Kruskal Wallis test with BH adjusted p-values. (C) The ordination plot (PCoA), with colours representing different protocols and shapes different preservation methods and with numbers representing the amount of stool used for the extraction (in mg).

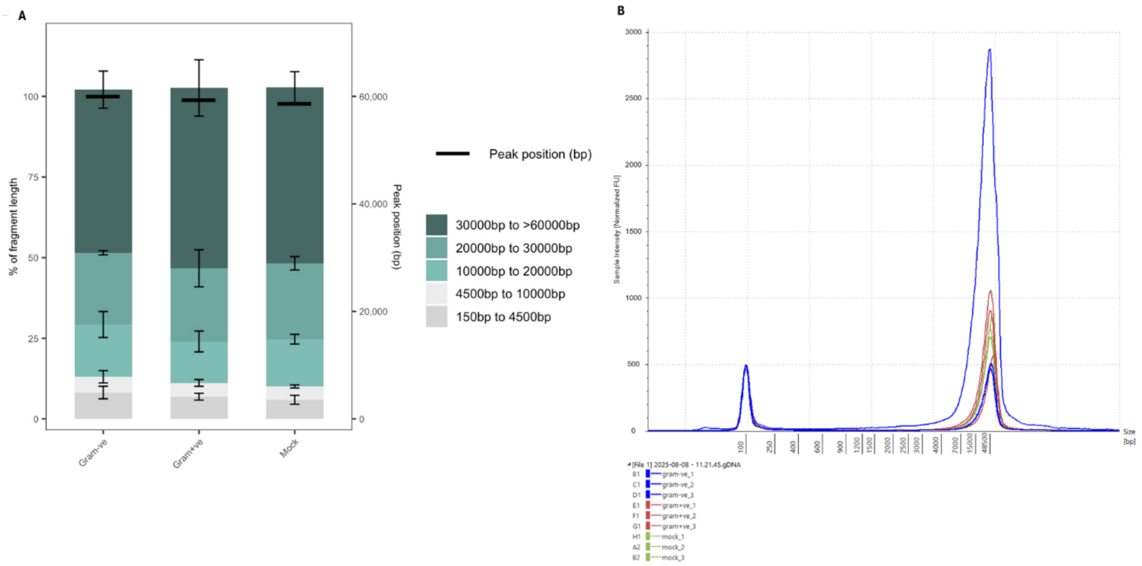


Supplement Fig.4: TapeStation fragmentation profiles of DNA of Single isolates and mock community samples extracted by Protocol 3 and preserved using DNA/RNA Shield (S). (A) The different user defined size classes of DNA fragmentation profiles plotted as stacked bars representing mean % of total DNA between replicates. Error bars represent SD between replicates and (B) TapeStation profiles of all samples/replicates.
